# Alpha-Synuclein co-pathology in Alzheimer’s Disease drives tau accumulation

**DOI:** 10.1101/2025.01.24.634706

**Authors:** Felix L. Struebing, Teodoro De Vecchi, Jeannine Widmann, Xiaoxuan Song, Federico Fierli, Viktoria Ruf, Qilin Tang, Thomas Arzberger, Sigrun Roeber, Thomas Koeglsperger, Otto Windl, Christian Haass, Juliane Winkelmann, Jochen Herms

## Abstract

The molecular basis for accelerated cognitive decline seen in Alzheimer’s Disease (AD) cases presenting with cortical alpha-Synuclein (α-Syn) co-pathology is not well understood. We show that such co-pathology brains express higher levels of microtubule-associated protein tau and that increasing α-Syn expression is sufficient to drive tau accumulation. Our results reveal a hitherto unknown link between the pathogenesis of AD and Parkinson’s Disease whereby tau and α-Syn synergistically drive dementia-related pathology.

## Main

Two of the most common neurodegenerative disorders, Alzheimer’s Disease (AD) and Parkinson’s Disease (PD), are classically characterized by pathological deposits of the microtubule-associated protein tau (tau) or alpha-Synuclein (α-Syn), respectively. However, both pathologies also co-occur in up to 50% of AD cases, suggesting a substantial cross-talk despite a different clinical presentation^1,2^.

The strongest molecular correlate to cognitive impairment, a unifying late symptom across most neurodegenerative disorders, is brain tau load^3^, yet recent studies suggest that tau and α-Syn can synergistically drive cognitive and functional decline^4,5^. This idea is corroborated by experiments demonstrating worse behavioral and cognitive outcomes in mouse and *in vitro* models of tau/α-Syn co-pathology^6–8^. However, a molecular explanation for this phenomenon is still lacking.

In this study, we re-evaluated neuropathologically diagnosed late-onset AD patients who had donated their brain tissue to the Neurobiobank Munich for α-Syn pathology as defined by McKeith^9^ and Braak^10^. Of the 136 cases, 35 had neocortical α-Syn pathology (hereafter as a group referred to as AD+ASYN), corresponding to the highest Braak stage 6, whereas 101 had no appreciable α-Syn pathology (hereafter termed AD, Figs. 1a and 1b). There was no difference in APOE genotypes between the AD and the AD+ASYN groups (p = 0.56), and they did not differ in the distribution of tau (Braak&Braak) stages^11^ (p = 0.15).

**Figure 1:**
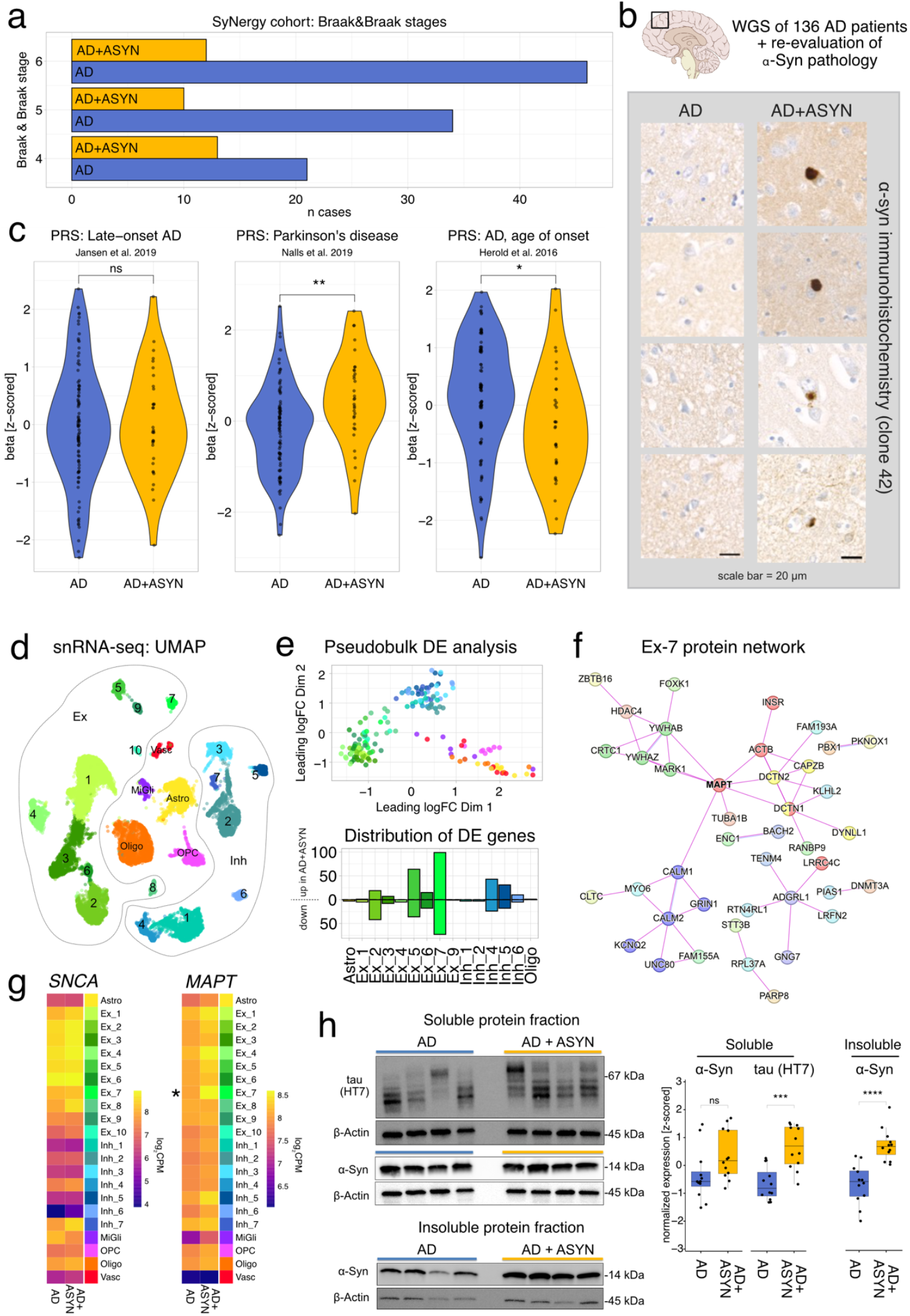
(a) Distribution of tau stages in the SyNergy whole-genome sequencing cohort. (b) Examples of α-Syn co-pathology in select samples. FFPE specimen were stained against alpha-synuclein (clone 42), and designated as AD+ASYN on presence of Lewy Bodies or Lewy neurites. (c) Polygenic risk scores for AD and AD+ASYN. (d) Dimensional reduction projection of single nucleus RNA-seq profiles. (e) Distribution of leading log2 fold-changes after pseudbulk aggregation and multidimensional scaling (top) and distribution of differentially expressed genes per cluster (bottom). (f) STRING protein-protein interaction network for the Ex-7 cluster using only experimentally validated interactions. (g) Heatmaps demonstrating cluster-wise *SNCA* and *MAPT* expression in AD and AD+ASYN brains. (h) Western Blots of select AD and AD+ASYN cases using antibodies against α-Syn (14H2L1), α-Syn phosphorylated at residue P129 (PS129), total tau (HT7), oligomeric tau (T22), and phosphorylated tau (AT8) normalized to ß-Actin.

To explore the genetic contributions related to α-Syn co-pathology, we performed whole genome sequencing and calculated polygenic risk scores (PRS, Fig. 1c). Using GWAS summary statistics from one of the largest AD studies^12^ did not reveal altered scores for late-onset AD between groups. However, there was a significant difference in age at onset for AD^13^ with a lower beta in the AD+ASYN group, suggesting that AD+ASYN individuals carry genetic risk factors associated with being affected by AD earlier in life. Importantly, we also found a statistically higher PRS for Parkinson’s disease^14^ in the AD+ASYN group, hinting at a role for PD-related risk factors in the manifestation of α-Syn co-pathology.

The most significant genetic risk factors for PD, rs356182 and rs356168, reside on chromosome 4q21, a locus that contains the gene encoding α-Syn (*SNCA*)^14,15^. We queried our whole genome sequencing data to test whether the higher PRS for PD in the AD+ASYN group could be explained by an enrichment of homozygous risk alleles within this region and found a suggestive association for rs356182 (p=0.08). This relationship became significant for rs356168 (p=0.04), where 25% of cases in the AD+ASYN group were homozygous for the risk allele, in contrast to only 10% of AD cases. The rs356168 risk variant was previously demonstrated to increase *SNCA* mRNA levels through the creation of new transcription factor binding sites^15^, thereby acting as expression quantitative trait locus.

Overexpression of *SNCA* represents a widely used animal and cell culture PD model, and additional *SNCA* copy numbers in humans are associated with familial PD^16,17^. To test whether AD+ASYN cases showed an increase in *SNCA* transcription, we performed single nucleus RNA-sequencing (snRNA-seq) of the superior frontal gyrus (SFG) in a subset (n=8) of AD and AD+ASYN cases that were matched in age and *APOE* genotype. Unsupervised clustering and annotation revealed the typical neocortical cell type composition, with large clusters of excitatory or inhibitory neurons, and smaller clusters of micro- or macroglia as well as vascular cells (Fig. 1d, Supp. Fig. 1).

We then performed differential expression testing using pseudobulk aggregates per group and cluster (Fig. 1e). In general, neuronal clusters had the highest number of differentially expressed genes (DEGs), with negligible numbers in the astrocyte (n = 3 genes) and oligodendrocyte (n = 1 gene) clusters (Supp. Table 1). The cluster with the highest number of DEGs was excitatory neuron cluster Ex-7. While many clusters, especially the excitatory ones, showed a slight increase in *SNCA* expression, this difference was never statistically significant (Fig. 1g), suggesting that neocortical α-Syn co-pathology is either not due to overexpression of its gene product, or due to it being spatiotemporally variable, rendering it invisible in post-mortem tissue.

We subsequently constructed cluster-specific protein networks from DEGs using only experimentally validated protein-protein interactions from the STRING database^18^ (Fig. 1f). For Ex-7, this revealed multiple subnetworks that surprisingly appeared to be interconnected by *MAPT*, the gene encoding microtubule-associated protein tau.

Indeed, *MAPT* transcripts were expressed at varyingly higher levels in all neuronal clusters, with a significant, almost two-fold upregulation in the AD+ASYN Ex-7 cluster (log_2_ fold-change: 0.89, Fig. 1g). We then tested SFG brain lysates from soluble and insoluble fractions for tau protein expression using antibodies against three different epitopes, together with antibodies against α-Syn (Fig. 1h, Supp. Fig. 2). As expected, the expression of α-Syn and α-Syn phosphorylated at residue S129 was significantly increased in the insoluble protein fraction from AD+ASYN brains, with a trend for increased α-Syn expression in the soluble fraction. We did not find higher levels of AT8, recognizing phosphorylated tau, or T22, specific for oligomeric tau, in AD+ASYN brains (Supp. Fig. 2). However, there was a significant increase in the expression of total tau (antibody HT7) in the soluble protein fraction of AD+ASYN brains, suggesting that α-Syn co-pathology is indeed characterized by higher levels of soluble, pre-pathogenic tau.

**Figure 2:**
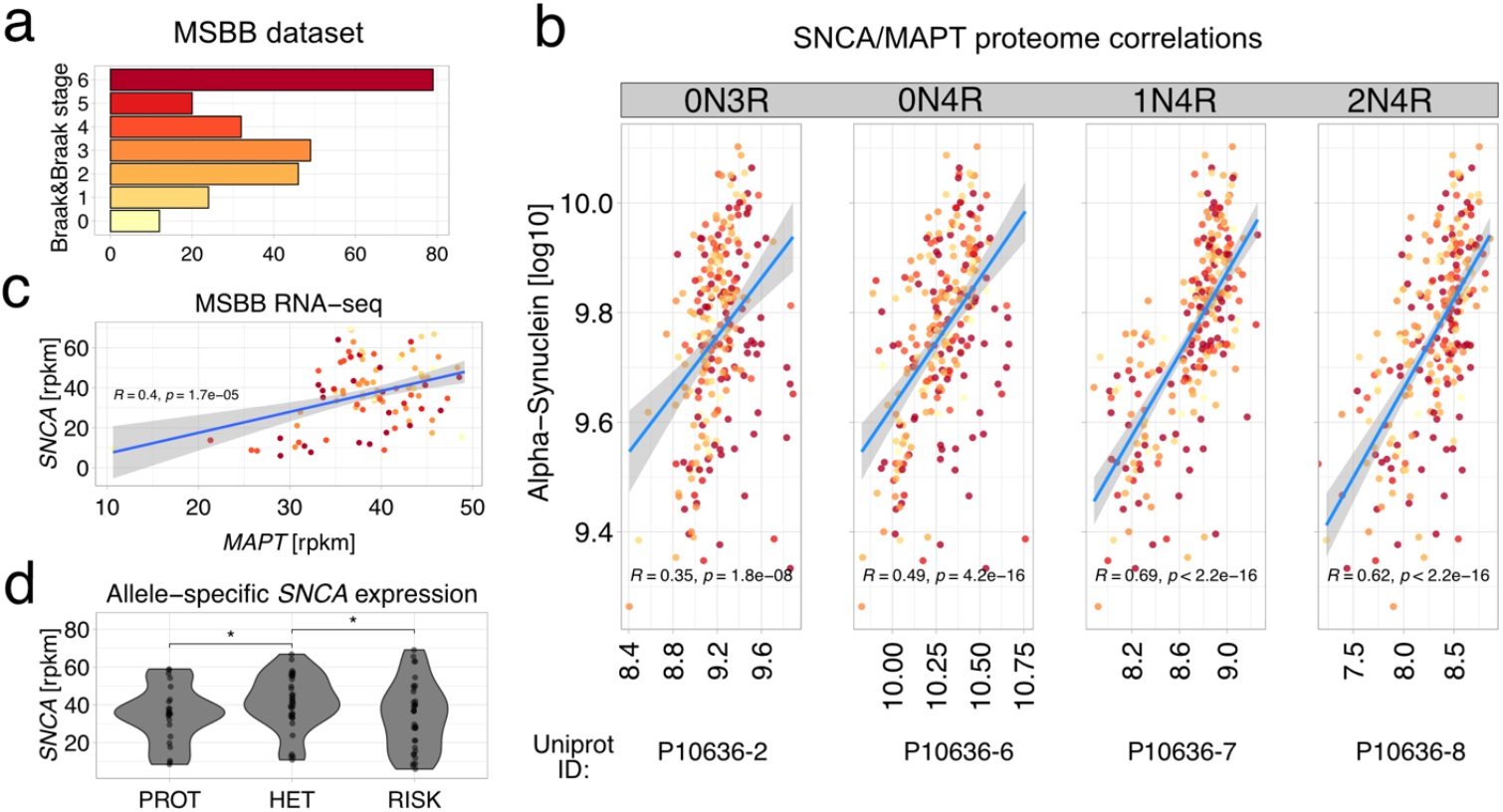
(a) Distribution of tau stages in the MSBB dataset with number of samples on the x-axis. (b) Correlation of α-Syn and tau protein in the MSBB proteomic dataset. Uniprot IDs for different tau isoforms is given on the bottom. (c) Correlation of *MAPT* and *SNCA* transcripts by RNA-seq. (d) Allele-specific expression for *SNCA* stratified by presence of rs356168. PROT = protective allele associated with decreased transcription. HET = heterozygote carriers. RISK = homozygous for the risk allele associated with increased *SNCA* transcription.

We then sought out to validate our findings in a larger cohort of post-mortem brain samples with varying stages of AD-related tau pathology, the Mount Sinai Brain Bank (MSBB) study^19^. This dataset provides neocortical proteome and transcriptome data from hundreds of replicates along with genotyping data (Fig. 2a). There was a strong, significant correlation in the protein expression of α-Syn and different tau isoforms (Fig. 2b). Likewise, we saw a positive relationship between *MAPT* and *SNCA* transcript counts in a subset of patients for whom neocortical RNA-seq data was available (R=0.4, p = 1.7e-5, Fig. 2c). Stratification by presence of rs356168, the *SNCA* allele previously found to be overrepresented in AD+ASYN brains of our whole-genome sequencing cohort, revealed higher *SNCA* expression for heterozygous and homozygous carriers of the risk allele (Fig. 2d).

To unequivocally define *SNCA* as a driver for *MAPT* expression, we overexpressed wildtype human *SNCA* in Lund Human Mesencephalic cells (LUHMES), a human dopaminergic cell line, by transduction with an adenovirus either carrying full-length human *SNCA* or GFP as control^20^ (Fig. 3a). Droplet digital PCR assays showed a significant increase in *MAPT* transcripts upon *SNCA* but not GFP expression (Fig. 3b). To exclude adenovirus-induced overexpression artefacts and better mirror the cell type environment found in human brains, we raised cerebral organoids^21^ from an iPS cell line either carrying four copies of the *SNCA* gene (AST, α-Syn triplication) or an isogenic control line in which a normal *SNCA* copy number had been restored (CAS, corrected α-Syn, Fig. 3c)^22^. Automated capillary Western blots of organoid lysates taken at different time points revealed a consistently higher expression of α-Syn in the AST line (Fig. 1d). After 70 days in culture, tau expression was also significantly increased in AST compared to CAS. Collectively, these data demonstrate a strong relationship between α-Syn and tau protein levels, a phenomenon at least partially driven by genetic risk factors associated with increased *SNCA* expression.

**Figure 3:**
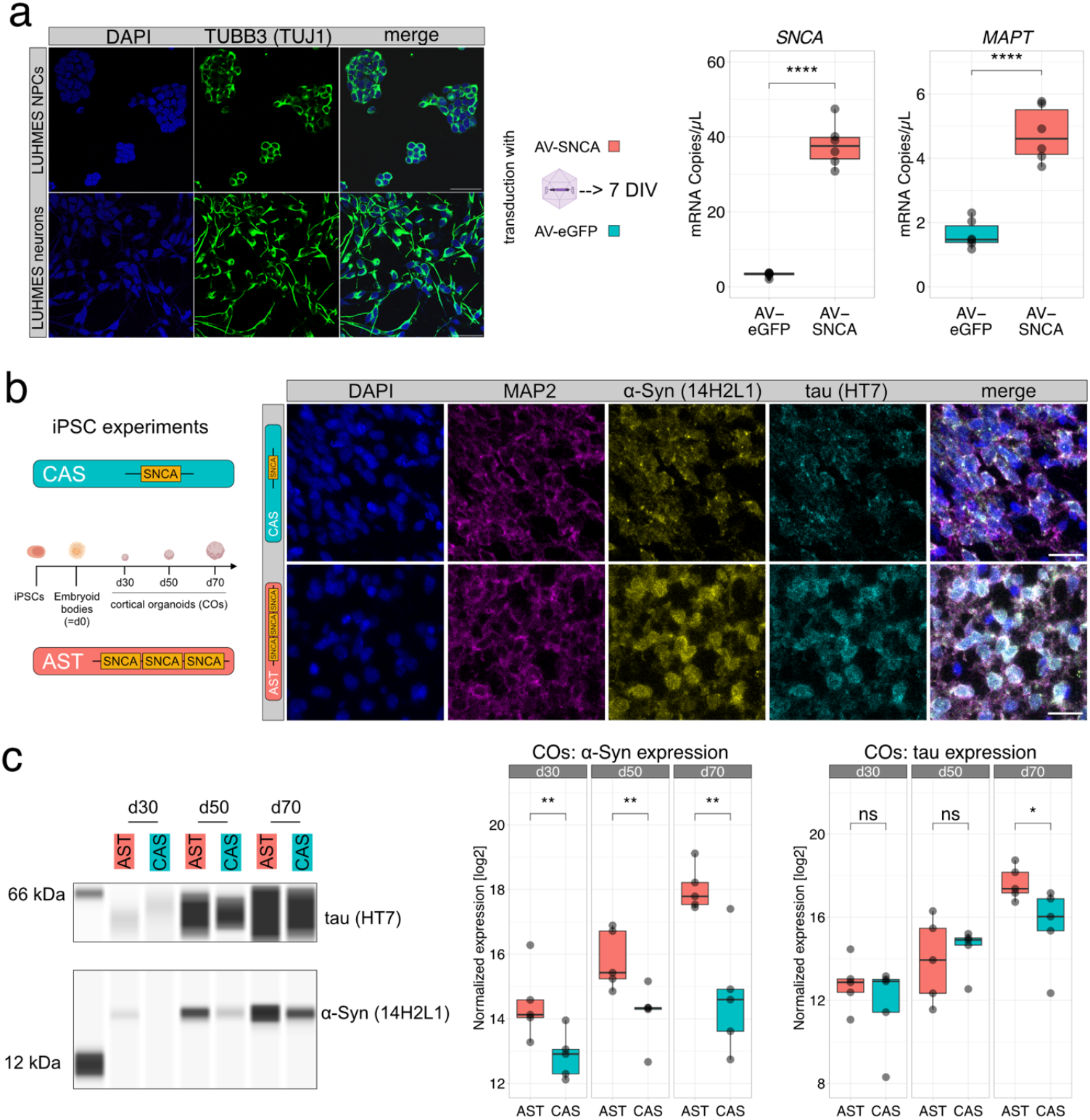
(a) Left: Immunofluorescence stainings for LUHMES progenitors (top) and differentiated LUHMES neurons. DAPI stains nuclei and TUJ1 beta-III-tubulin. Right: Absolute concentration of SNCA and MAPT transcripts after transduction with adenovirus carrying full-length SNCA or eGFP as control measured by digital droplet PCR. (b) Immunofluorescent images of cortical organoids after 70 days in culture. (c) Tau and α-Syn protein expression (automated capillary electrophoresis / SimpleWestern™) normalized to total protein content of cerebral organoids at different timepoints demonstrating higher α-Syn expression in the AST line throughout all time points and higher tau expression after 70 days in culture.

The simultaneous presence of tau and α-Syn pathologies in AD, the classic form of which is only characterized by tau and ß-Amyloid plaques, has puzzled the field for almost 40 years^23^. Previous large-scale studies have demonstrated earlier disease onset and earlier death for subjects with α-Syn co-pathology in AD^24^, a finding to which we can now add the insight that this phenomenon is likely due to altered genetic risk scores and thus heritable. Mouse studies have laid out a foundation for a putative mechanism: In the absence of tau, α-Syn spreading was reduced, but not the other way around, suggesting that an increased amount of α-Syn accelerates the disease phenotype^7^. Our data bring these findings to the smallest common denominator, suggesting that increased *SNCA* transcription also leads to upregulation of its protein product α-Syn, and that this upregulation is sufficient to drive accumulation of tau, also by upregulating its transcripts from the *MAPT* locus.

In conclusion, our data demonstrate a role for α-Syn co-pathology in driving tau accumulation in AD, and because tau load is strongly correlated to cognitive decline, they offer an elegant mechanistic explanation for a finding already noted by clinicians and neuropathologists decades ago. Future experiments should investigate the genomic networks kickstarted by *SNCA* transcription to define targetable pathways or bring us at least one step closer to a comprehensive understanding of the complex molecular processes taking part during neurodegeneration.

## Online Methods

### Human cohort and neuropathological assessment

All participants included in the study had given informed consent to donate their brain according to the Code of Conduct laid out by the BrainNet Europe^25^. At autopsy, brain hemispheres were treated differently: The left hemisphere was fixed in formalin for a duration of two weeks or longer, while the right hemisphere was snap-frozen immediately. From the former, paraffin-embedded specimen sampled across the whole cerebrum, brain stem, cerebellum, and spinal cord were used for diagnostic examination. Tau staging was performed according to Braak&Braak^11^ by staining appropriate areas with the monoclonal antibody AT8 (ThermoFisher, #MN1020). Alpha-Synuclein pathology was assessed as defined by Braak^10^ and McKeith^9^ using clone 42 (abcam, ab280377). All specimens were evaluated by at least two board-certified neuropathologists and only samples neuropathologically diagnosed with AD entered the study. Importantly, subjects with a neuropathological diagnosis of Lewy Body Disease (DLB/PDD), significant co-pathology besides α-Syn co-pathology, or unclear cases were excluded from the study. AD+ASYN brains included samples with neocortical Lewy Body or Lewy Neurite pathology (Fig. 1b). Sample information for WGS can be found in Supplemental Table 3.

### Whole genome sequencing, variant calling and polygenic risk scores

DNA was isolated from 1 cm^3^ large tissue cubes taken from fresh-frozen cerebellum using the QIAmp DNA Mini Kit (Qiagen, 51304). Library preparation was performed with the TruSeq PCR-free genomic DNA library prep kit (Illumina, FC-121-3003) according to the manufacturer’s instructions. Libraries underwent 2×150 bp paired-end sequencing on an Illumina NovaSeq machine until a minimum depth of 35X was reached. Alignment and variant calling were performed using a Snakemake pipeline incorporating the GATK best practices. Briefly, after FastQC and adapter trimming, alignment to the hs1/T2T genome assembly (chm13v2.0) was performed with BWA-MEM2. Variant calling, recalibration and joint genotyping were done using GATK version 4.0^26^. Ultimately, samples with familial AD or PD mutations (PSEN1/2, APP, SNCA, MAPT) were excluded from the study. Polygenic risk scores were calculated with PRSKB^27^ by supplying GWAS summary statistics from relevant studies.

### Single-nucleus RNA sequencing

Fresh-frozen human cortical tissue was microdissected, homogenized in NP40 buffer and filtered through a 70 μm strainer. After incubation on ice and centrifugation, nuclear pellets were resuspended in PBS with 1% BSA and RNase inhibitor, and nuclei were stained with 7AAD before being sorted into BSA containing RNase inhibitor using a Sony SH800 cell sorter. GEMs were generated on a 10x Chromium controller with a targeted nuclei number of ∼4000 per sample to minimize doublet generation. RNA library construction was performed with a 3’ gene expression kit (10X Genomics, PN-1000285) according to the manufacturer’s instructions. Following verification of correct insert sizes using a BioAnalyzer with the DNA High Sensitivity Chip (Agilent, 5067-4626), molarities were determined by droplet digital PCR (see below), and libraries were pooled in an equimolar fashion. Sequencing took place on an Illumina NovaSeq using an S2 flow cell with 2×150 cycles.

### Bioinformatic and statistical analysis

Single nucleus RNA libraries were aligned to the hs1/T2T genome (chm13v2.0) using STARSolo^28^ while accounting for each sample’s variants by supplying sample-specific vcf files to the --varVCFfile argument. After QC, which included removing nuclei with small coverage or a large fraction of mitochondrial reads, we retained ∼27,000 nuclei for downstream analysis. Each sample was normalized separately with the SCTransform v2 algorithm provided by the Seurat R package to account for differing library sizes^29^.

Normalized samples were merged into one Seurat object, on which PCA and unsupervised clustering were carried out using the SNN algorithm on the top 50 principal components. Optimal cluster numbers and clustering stability were assessed by ClustAssess^30^. Top-level cluster annotations were retrieved by referencing our data set with CELLxGENE^31^, a transformer-model-based, single cell sequencing-derived human cell type compendium, yielding excitatory, inhibitory, glial and vascular clusters. These clusters were further refined for glial cells by referencing known cell type markers for OPCs, astrocytes, microglia and oligodendrocytes, respectively, whereas neuronal clusters were numbered in ascending order according to their size. For visualization purposes, we removed batch effects by integrating samples with Harmony^32^, followed by dimensional reduction with UMAP. Pseudobulk testing was performed by aggregating gene counts by cluster and group, removing genes with low coverage, fitting robust negative binomial generalized linear models and conducting quasi-likelihood tests using the edgeR package^33^. Nominal P-values were corrected for multiple comparisons using the Benjamini-Hochberg procedure. We considered genes with an absolute log2-fold-change of > 0.5 and an FDR < 0.1 as statistically significant. STRING network construction was done by uploading DE genes per cluster to STRINGdb and filtering for experimentally validated protein-protein interactions. Computational analyses were performed on an HPC running Arch Linux and the R statistical programming environment version 4.3.3. Data processing pipelines and plots throughout the manuscript made heavy use of the tidyverse and ggplot2 packages for R. Significance levels are denoted by asterisks: * = p < 0.05; ** = p < 0.01; *** = p < 0.001; **** = p < 0.0001.

### External data analysis

The results published here are in whole or in part based on data obtained from the AD Knowledge Portal (https://adknowledgeportal.org/). These data were generated from postmortem brain tissue collected through the Mount Sinai VA Medical Center Brain Bank and were provided by Dr. Eric Schadt from Mount Sinai School of Medicine, and proteome data were provided by Dr. Levey from Emory University. After receiving the appropriate permissions, MSBB data was downloaded from www.synapse.org. Normalized proteome data was available for n = 310 individuals and ∼90,000 Uniprot IDs, the gene symbols of which were retrieved by querying Uniprot’s REST API. RNA-seq data was downloaded for a subset of these cases for which both RNA counts and genotyping information (as a vcf file) was available (n = 107), and cases were joined on their unique individual identifier. Genotype and transcriptome data came aligned to the hg19 genome assembly, and featureCounts from the Rsubread package was used for assigning counts to genes with default arguments.

### Protein isolation and Western blotting

For automated capillary Western blots, homogenized iPSC-derived cerebral organoids were lysed in 1x RIPA Lysis Buffer (Merck, #20-188), and the protein levels of the homogenates were quantified by bicinchoninic acid assays (Sigma-Aldrich). Protein samples were reduced in Fluorescent Master Mix with 200 mM dithiothreitol (ProteinSimple) and run on a Jess Automated Western Blot System (ProteinSimple) using the 12-230 kDa Separation Module and 25-Capillary Cartridges (ProteinSimple). Proteins in the capillaries were blocked using Antibody Diluent 2 (ProteinSimple) and incubated with corresponding primary antibodies (anti-alpha Synuclein (14H2L1) (Invitrogen, 1:50), anti-Tau (HT7) (Invitrogen, 1:50) for 90 minutes at room temperature and then with secondary HRP-conjugated antibodies (anti-rabbit and anti-mouse (ProteinSimple)) for 30 minutes at room temperature. For chemiluminescence detection, proteins were incubated in Luminol-Peroxide Mix (ProteinSimple) and scanned by the Jess Automated Western Blot System (ProteinSimple).

For regular Western blots from post-mortem brain tissue, samples were homogenized in 1x RIPA lysis buffer (Merck, #20-188) supplemented with protease inhibitors (Roche, CO-RO) and phosphatase inhibitors (Roche, PHOSS-RO) using a Mini Bead Mill Homogenizer (VWR). After 30 min incubation on ice, the lysates were centrifuged at 16,000g for 30 min at 4°C to obtain the soluble protein fraction. The remaining pellets were washed once with lysis buffer and re-homogenized in 1% sarkosyl-containing lysis buffer, rotated overnight at 4°C. The samples were again centrifuged at 16,000g for 30 min at 4°C and the supernatant was collected as insoluble protein fraction. The protein concentration was determined by the Pierce™ BCA Protein Assay (Thermo Scientific, #23225). 15μg of proteins for each sample were run on a 4-20% precast polyacrylamide gel (Bio-Rad, #4561094) and electrophoresis was run at 90V for 1h and then 120V for 1h. The proteins were transferred to 0.45μm PVDF membranes (Millipore, #IPVH00010) at 200mA for 2h. The membranes were post-fixed with 4% paraformaldehyde and 0.1% glutaraldehyde for 30 min, washed in TBS supplemented with 0.1% Tween-20 (Sigma-Aldrich, #93773, TBST) and then blocked with 5% BSA (Sigma-Aldrich, #A7030) in TBST for 1h at room temperature (RT). The membranes were probed with primary antibodies diluted in the blocking solution overnight at 4°C. The next day, the membranes were washed 3 times with TBST and incubated with secondary antibodies for 1h at RT. After washing 3 times, the blots were developed with Pierce™ ECL Western Blotting Substrate (Thermo Scientific, #32109) and imaged on the ChemiDoc MP Imaging System (Bio-Rad). For stripping, the blots were washed 3 times with TBST and then incubated in stripping buffer (ThermoFisher, #46430) for 15 minutes at RT. Thereafter, the blots were washed and re-probed as before. The following primary and secondary antibodies were used: HT7 anti-Tau (ThermoFisher, #MN1000), AT8 anti-phosphorylated Tau (ThermoFisher, #MN1020), T22 anti-Tau oligomer (Sigma-Aldrich, #ABN454), 14H2L1 anti-alpha synuclein (ThermoFisher, #701085), PS129 anti-phosphorylated alpha synuclein (Abcam, #ab51253), anti-beta actin (ThermoFisher, #AM4302), anti-mouse IgG HRP-conjugate (Sigma-Aldrich, #12-349), anti-rabbit IgG HRP-conjugate (Promega, #W4011). Quantification of band densities was analyzed by FIJI software. Each target protein was normalized to beta actin. A two-tailed unpaired t-test was performed to analyze the z-scored data.

### RNA isolation and digital droplet PCR

Single-nucleus RNA-seq libraries were quantified using the ddPCR Library Quantification Kit for Illumina TruSeq (Bio-Rad, #186-3040) according to the manufacturer’s instructions. For gene expression assays, total RNA was isolated with the RNeasy Mini Kit (Qiagen, #74104) and quantified by fluorometry using the Qubit dsDNA HS assay (ThermoFisher, Q33230) before 1 ng of RNA were retrotranscribed with the QuantiTect Reverse Transcription Kit (Qiagen, #205311). After droplet generation with 1 μL of RT reaction, 10 μL of 2X ddPCR EvaGreen Supermix (Bio-Rad, #1864033), 1 μL of each 10 μM forward and reverse primer and 7 μL of H2O, PCR was performed with a Tm of 60C on a Bio-Rad C1000 Touch Thermal Cycler. Ultimately, droplets were read on a QX200 droplet reader (Bio-Rad). Primer sequences are given in Supp. Table 2.

### LUHMES cell culture experiments

LUHMES cells were seeded into appropriate culture vessels pre-coated with 1% of PLO 100x (10 mg/ml) (Sigma-Aldrich, P3655) and 99% of DPBS without calcium and magnesium (Thermo-Fisher, 14190144). Cell culture media for LUHMES was composed of DMEM/F12 (Thermo-Fisher, 11330032), 1% N2 supplement 100x (Thermo-Fisher, 17502001) and bFGF (25 μg/ml) (Thermo-Fisher, 11330032). For differentiation, the PLO solution was removed from the culture vessel, washed 3 times with DPBS, and a solution of 0,5% Fibronectin (5 μg/ml) (Sigma-Aldrich, F0895) in UltraPure DNase/RNase-Free Distilled Water (Invitrogen, 10977023). 100 ml of the medium used for differentiation into post-mitotic neuronal cells were composed of 98 ml of DMEM/F12, 1 ml of N2 supplement 100x, 1 ml of dibutryl cAMP (49 mg/ml) (Sigma-Aldrich, D0627), 40 μl of glial cell derived neurotrophic factor (GNDF) (5 ng/μl) (R&D System, 212-GD-010) and 100 μl of Tetracycline (1 mg/ml) (Sigma-Aldrich, T7660). To split 70% to 80% confluent cells, they were washed once with DPBS and afterwards incubated for 5 minutes at 37°C with Trypsin-EDTA 0,05%. Trypsinization was blocked by adding an equivalent volume of a solution made with 10% FCS or FBS and 90% DMEM-F12. Trypan-Blue-stained cells were counted using a Neubauer counting chamber. For differentiation with a density of 1 million cells per well of a 6-well-plate, cells were left growing for 3 days in a T75 flask in order to arrive at confluence. 100 μl of a solution composed of differentiation medium and an adenovirus expressing either human wild-type SNCA or eGFP under the control of a CMV promoter was added to each well at a multiplicity of infection of 1.5 at day 2 after initiation of the differentiation process. After 24 hours, the cells were washed once with DPBS and medium was replaced with 2 ml of differentiation medium per well.

### iPSC culture and cerebral organoids

Induced pluripotent stem cell (iPSC) lines were a gift from Tilo Kunath, University of Edinburgh. The first iPS cell line used in this study was derived from fibroblasts of an Iowan family with early-onset PD harbouring a *SNCA* triplication^17^ and is herein referred as AST (α-Synuclein triplication iPS cell line). The second line used is an isogenic, CRISPR-Cas9 corrected iPSC line with a normal *SNCA* copy number, that has been generated from the AST cell line and is herein referred to as CAS^34^ (corrected α-Syn triplication iPS cell line).

Both the AST and CAS iPSC lines and subsequently differentiated cells were maintained in an incubator at 37 °C with 5% CO_2_ throughout the experiments. Cell culture vessels for iPSC culture were coated with Matrigel (Corning) diluted 1:100 in DMEM/F12 (Thermo Fischer) The Matrigel-DMEM/F12 solution was applied to the culture vessels covering the surface and the plates were incubated for 1h at 37 °C. Before thawing and plating the cells, the culture vessels were washed three times with DPBS -/- (Thermo Fisher). mTesR+ medium (STEMCELL Technologies) with 10 μM Rock-Inhibitor (Merck) was added in each well and incubated at 37 °C. Frozen iPSCs were thawed in a 37 °C water bath until the ice block detached from the tube. The cells were decanted into a 15 ml centrifuge tube with 10 ml pre-warmed DMEM/F12 and the empty cryovial was washed with 1 ml DMEM/F12. The 15 ml tube was gently tilted to mix the cells and subsequently pelleted at 300 x g for 5 minutes at RT. The supernatant was removed and the pellet was carefully resuspended in 0.5 ml mTesR+ medium per well. The cell suspension was applied onto the plate, carefully moved with quick side movements to evenly distribute the cells and placed in the incubator. After 24 hours, the medium was changed to remove cellular debris and Rock-Inhibitor. Until expansion of the cells, the medium was changed every second day until the cells reached confluency for passaging.

Cells were passaged on prepared multi-well plates coated with Matrigel diluted 1:100 in DMEM/F12 for 1 h at 37 °C. For routine cell passaging, iPSCs being 70-80% confluent were passaged as colonies using ReLeSR (STEMCELL Technologies). The consumed medium was aspirated, replaced with 1 ml ReLeSR reagent and incubated for 1 minute at room temperature. ReLeSR was removed and the plate was placed in the incubator for 5 minutes. Cell colonies were detached by firmly tapping the side of the well plate and collected in 2 ml DMEM/F12 medium, centrifuged for 5 minutes at 300 x g at RT, resuspended in the desired mTesR+ volume and evenly distributed with quick side movements. Cells were routinely passaged in ratios between 1:3 and 1:6. When starting experiments with a pre-defined number of cells, iPS cells were harvested using Gentle Cell Dissociation Reagent (GCDR) (STEMCELL Technologies) to obtain single cells. The consumed medium was aspirated and replaced with 1 ml GCDR and incubated for 5-8 minutes at RT. After incubation, 1 ml of DMEM/F12 was added to the wells and the cells were scratched off using a cell scraper. The cells were collected and counted using a Neubauer counting chamber system, centrifuged for 5 minutes at 300 x g and eventually plated at the desired density. Being re-plated as single cells, Rock-Inhibitor at 10 μM final concentration was added to the culture medium for 24 hours. Cerebral organoids (CO) were cultured following the Lancaster Protocol^21^ using the Cerebral Organoid Kit and Cerebral Organoid Maturation Kit (STEMCELL Technologies) for subsequent maturation of the organoids. On day 0, CO generation was started with embryoid body (EB) formation. Therefore, one 70-80% confluent well of a 6-well plate of iPSC cells was washed with PBS and harvested with Gentle Cell Dissociation Reagent (STEMCELL Technologies). Cells were resuspended in EB seeding medium and 100 μl of cell suspension were plated into each well of a 96-well round-bottom ultra-low attachment plate (Corning) with a concentration of 90.000 cells/well. On day 2 and 4, 100 μl EB formation medium was added per well. On day 5, small formed EBs were transferred with a wide-bore pipette tip onto an ultra-low attachment 24-well plate (Corning) with 500 μl of fresh induction medium. At day 7, the EBs were collected with a wide-bore pipette tip and placed on an embedding sheet (STEMCELL Technologies). Excess medium was removed and a 15 μl Matrigel (Corning) droplet was added to the EB, which was subsequently positioned in the center of the drop. After 30 minutes incubation at 37 °C for Matrigel polymerization, the Matrigel-embedded EBs were washed off the embedding sheet with expansion medium into an ultra-low attachment 6-well plate (Corning) and incubated for 3 days. At day 10, the consumed medium was replaced with fresh maturation medium and the well plate was transferred to an orbital shaker at 75 rpm at 37 °C. Media changes were performed routinely every 2-3 days. Organoids ready for harvesting were taken from the plate using a 1000 μl wide bore pipette tip. PCR Mycoplasma tests (Promokine) were performed routinely to exclude cell culture contamination.

### Immunofluorescence

To stain LUHMES cells, a coated coverslip was placed into a well of 24-well plate and cells were added (0.2 × 10^6^ for LUHMES and 0.5 × 10^6^ for differentiated neuronal cells). The coverslip was moved to another plate and washed once with DPBS. Afterwards, it was incubated for 15 minutes with 4% paraformaldehyde (PFA) and washed 3 times for 10 minutes each with PBS. Incubation followed with 200 μl of a 1:200 primary antibody dilution (Anti βIII-Tubulin Rabbit) with 0.05% Tween 20 and 2% I-Block reagent (Sigma). The coverslip was incubated at 4°C overnight and then washed 3 times with PBS. 200 μl of a secondary antibody (Anti-Rabbit Alexa 488) and DAPI (both in 1 1:1000 dilution) were added and the coverslip was incubated for 2 hours at room temperature. After 3 washes in DPBS, each lasting 5 minutes, the coverslip was mounted on microscopy slides using mounting medium (Histo-Line laboratories, PMT030) and sealed with nail polish.

Cortical organoids were fixed in 4% PFA, embedded in paraffin and cut to 10 μm thick sections. For deparaffinization, the slides were immersed in Xylene twice for 10 min, 100% Ethanol twice for 10 min, 95% Ethanol for 5 min, 70% Ethanol for 5 min, 50% Ethanol for 5 min, followed by a quick rinse with deionized H2O and incubation in PBS for 10 min. Antigen retrieval was performed by immersing the slides in 1x Sodium Citrate solution (Abcam, #ab64214) and placing them in a pressure cooker at high pressure for 20 min. The sections were cooled to RT and washed in PBS 3 times for 5 min. For permeabilization, the slides were then incubated with 0.2% Triton X-100 (Sigma-Aldrich, #T9284) in PBS for 10 min. After washing 3 times with PBS, the slides were blocked with 0.3% Triton X-100 in PBS with 5% normal goat serum (Sigma-Aldrich, #G9023) for 2h at RT. Primary antibodies were diluted in blocking solution and incubated overnight at 4°C. The next day, slides were washed 3 times with PBS and then incubated with diluted secondary antibodies in blocking solution for 2h at RT. The following primary and secondary antibodies were used: HT7 anti-Tau (ThermoFisher, #MN1000), 14H2L1 anti-alpha synuclein (ThermoFisher, #701085), anti-MAP2 (SYSY antibodies, #188004), Alexa 647 (ThermoFisher, #A32787), Alexa 555 (ThermoFisher, #A21429) and Alexa 488 (ThermoFisher, #A11073). The sections were washed 3 times with PBS and covered with ROTI® Mount FluorCare DAPI (ROTH, #HP20.1). All images were acquired using the confocal microscope Stellaris 5 (Leica). For each immunofluorescence reaction, secondary-only controls were verified to be absent of fluorescent signal before proceeding. Laser power and detector gain levels remained unchanged between imaging sessions across groups.

## Data and Code availability

All code used in this study is available under https://github.com/fstrueb/MAPT_SNCA. Human WGS and snRNA-seq data can be shared upon request after appropriate review by the LMU Ethics committee. Access to MSBB data can be gained after institutional review, see www.synapse.org.

## Acknowledgements

Human induced pluripotent stem cells were a gift from Dr. Tilo Kunath (University of Edinburgh, UK). We would like to acknowledge Vanessa Boll for DNA isolation, Dr. Norbert Buresch for dissection of fresh-frozen brain tissue, and Michael Schmidt for his help with immunohistochemistry and slide scanning. We would also like to thank Dr. Oliver Keppler for granting us access to the SimpleWestern™ apparatus. Whole genomes were sequenced by the Helmholtz Munich Core Facility Genomics (lead by Dr. Inti Alberto de la Rosa Velazquez) and generously sponsored by the Munich SyNergy consortium. We are thankful to Dr. Paul Feyen, Dr. Lars Paeger, Dr. Nils Briel, Antonia Neubauer and Dr. Patrick Harter for inspiring discussions. Ultimately, we are deeply indebted to the hundreds of patients who chose to donate their brains to research, including their families.

## Author contributions

FLS and JH conceived the project. JH, VR, TA and SR evaluated brains of the WGS cohort for alpha-synuclein pathology. JuWi, CH and JH organized and oversaw WGS. OW supplied material from the Munich Brain Bank. FLS generated snRNA-seq data for suitable cases as identified by VR. FLS performed WGS variant calling and all bioinformatic and statistical analyses. Digital droplet PCR assays were run by FLS, TDV, JeaWi and XXS. JeaWi raised cortical organoids and performed Mycoplasma testing. TDV performed all SimpleWestern™ experiments and assisted in growing cortical organoids. FF, QLT and TK ran LUHMES experiments. XXS performed Western blots of post-mortem tissue and immunofluorescent stainings of organoids. FLS wrote the first draft of the manuscript and designed figures with significant input from JH, TDV and XXS. All authors read, improved and approved the final manuscript.

