## Supplemental Figures for "Alpha-Synuclein co-pathology in Alzheimer’s Disease drives tau accumulation"

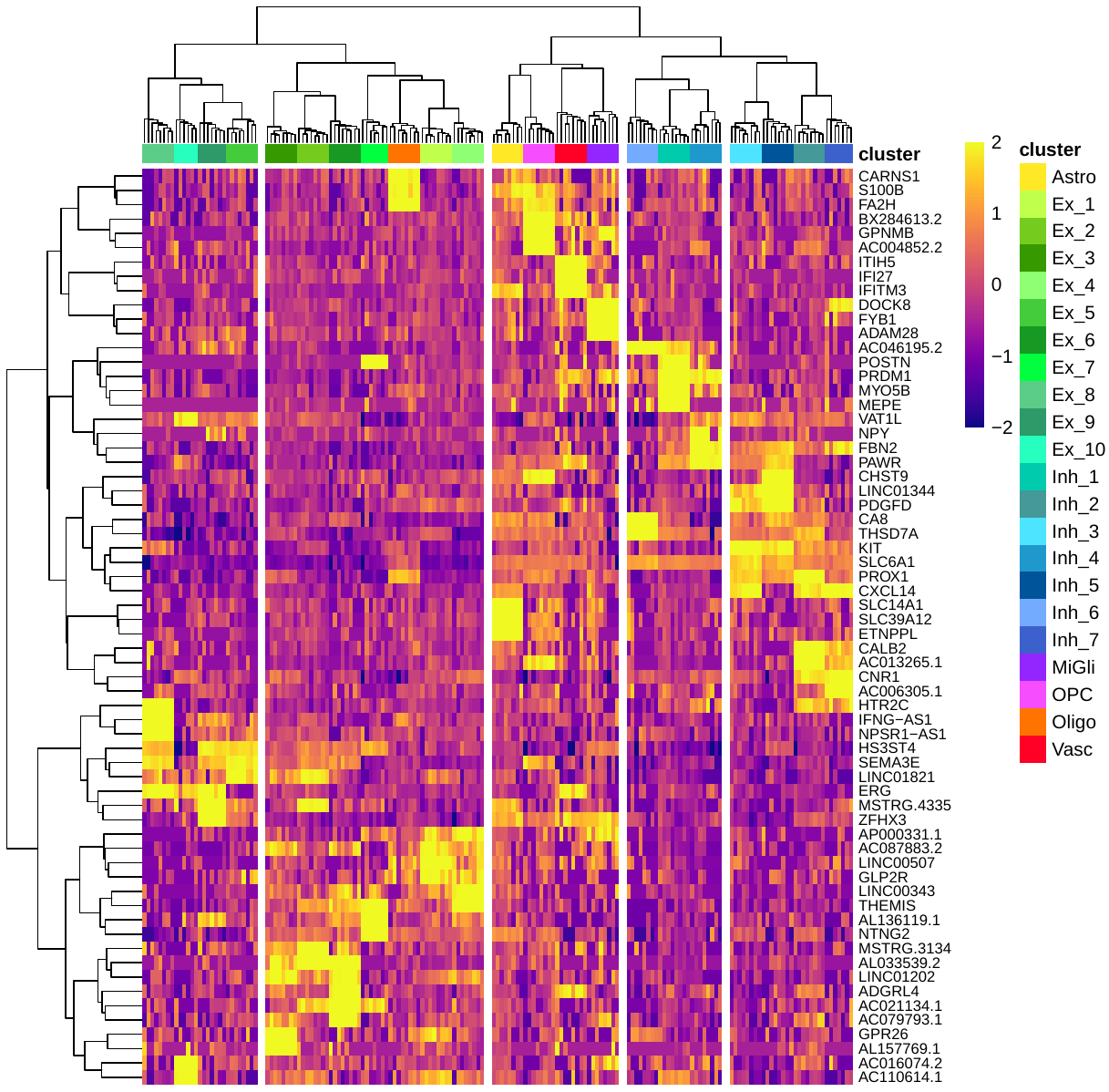

Supp. Fig. 1: Marker heatmap for snRNA-seq clusters. The top 3 identifying genes per cluster are shown. The scale corresponds to z-scored expression values.

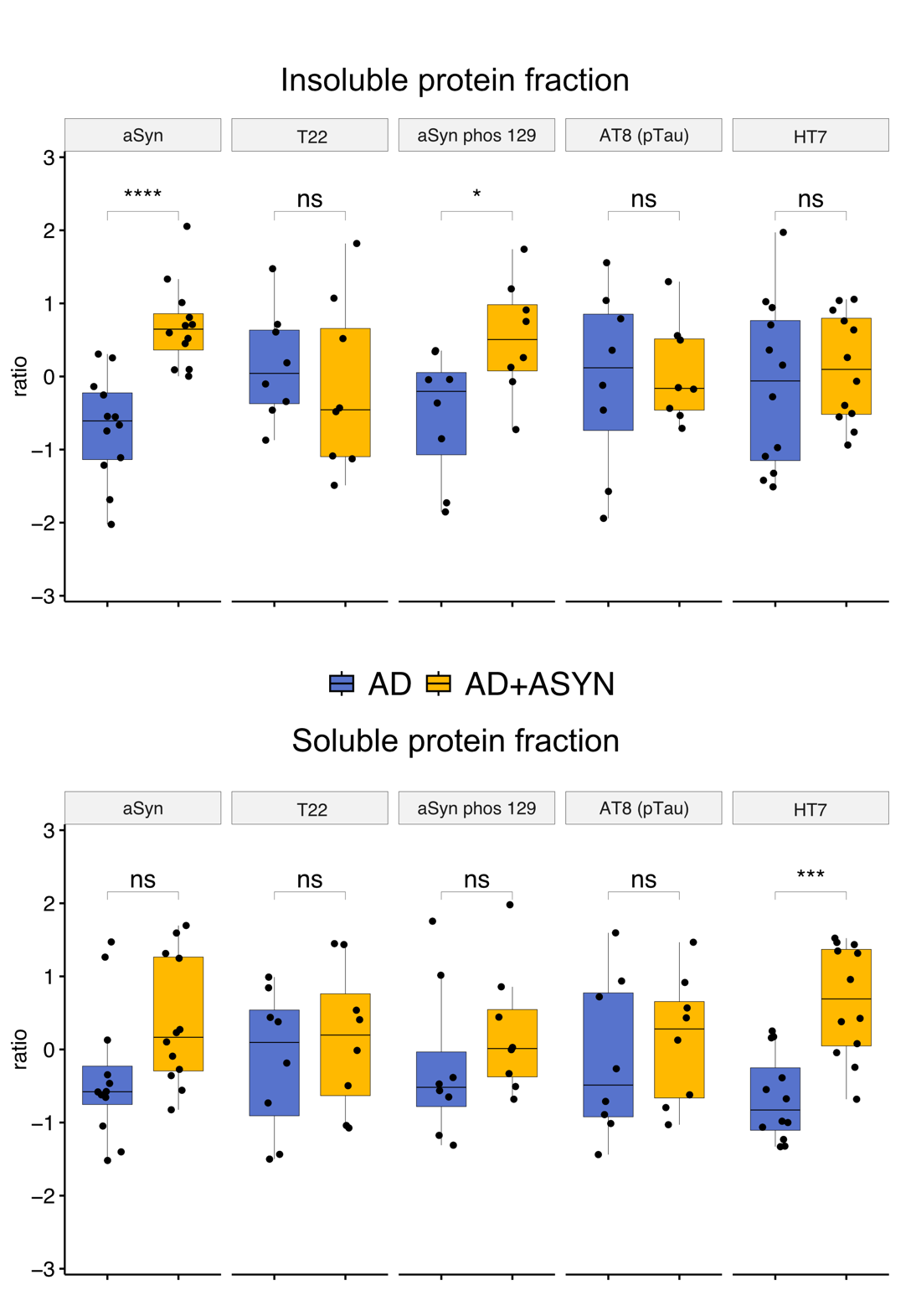

Supp. Fig. 2: Quantifications of Western Blots from post-mortem human neocortical tissue against different epitopes of alpha-synuclein and tau.
